# Predicting stress response trajectories: Differential contributions of limbic and prefrontal regions to cortisol and affective responses

**DOI:** 10.1101/2025.11.14.688580

**Authors:** Renée Lipka, Ludwig Kreuzpointner, Christoph Bärtl, Marina Giglberger, Julian Konzok, Hannah L. Peter, Nina Speicher, Lea Waller, Brigitte M. Kudielka, Stefan Wüst, Henrik Walter, Gina-Isabelle Henze

## Abstract

Why do individuals respond differently to stress? Since rodent studies indicated that stress regulation relies on limbic and medial prefrontal cortex (mPFC) outputs, we aimed to investigate whether data from these regions could also predict cortisol and affect trajectories following psychosocial stress in humans. In this pre-registered study, 281 healthy adults (145 female) were exposed to Scan*STRESS*. Repeated assessments of salivary cortisol and negative affect were used to identify response trajectories (i.e. groups of participants) using latent class mixture modelling (LCMM). LCMMs without brain predictors were compared to LCMMs including structural (volume, thickness) and functional (activation, exposure-time effect) predictors from the amygdala, hippocampus, or mPFC regions. Results showed that cortisol LCMMs without brain predictors exhibited a single mean trajectory. Adding brain predictors resulted in three to four response trajectories, depending on region and outcome. Within identified models, cortisol ‘hyper-response’ trajectories were predicted by larger amygdala and hippocampus volumes. Cortisol ‘non-responses’ were predicted by greater amygdala activation and volume. ‘Elevated baseline’ cortisol was predicted by higher hippocampal activation. mPFC markers did not predict cortisol trajectories, however, medial orbitofrontal cortex parameters identified affect response profiles mirroring trait-like affect. Together, our findings suggest dissociated roles of limbic and mPFC regions in stress regulation: While limbic structures predicted cortisol responses, the mPFC shaped affective experience.

## Introduction

People respond differently to stress. This variability is evident in subjective emotional experience as well as cortisol output, the end-product of the hypothalamic-pituitary-adrenal (HPA) axis. Following exposure to a stressor, the HPA axis becomes activated with the hypothalamus secreting corticotropin-releasing hormone (CRH), which in turn stimulates adrenocorticotropic hormone (ACTH) release from the pituitary, finally resulting in cortisol secretion from the adrenal glands. The temporal dynamics of this response determine their effect: During the first 20 minutes, cortisol primarily binds to high-affinity mineralocorticoid receptors (MRs), which facilitates risk assessment, memory retrieval, and response selection. Subsequently, rising cortisol levels also engage lower-affinity glucocorticoid receptors (GRs), which promotes memory consolidation and negative feedback terminating HPA axis activity (1,2). If an individual copes successfully, rapid rises and recovery of cortisol serve as windows of adaptive plasticity, forming internal schemes of control and safety promoting resilience (2). In contrast, deviations from the normative response, such as exaggerated, prolonged, or flattened cortisol responses have been associated with a range of stress-related physical and mental health outcomes, including depression, anxiety, chronic pain, and fatigue (3,4). Insights into cortisol variability therefore have implications for both prevention and treatment interventions.

Traditionally, variability in cortisol stress responsivity has been investigated by grouping participants into responders or non-responders, or by using summary metrics (e.g. difference scores, areas under the curve) (5,6), which mask temporal dynamics. Latent class mixture modelling (LCMM) overcomes this limitation by grouping individuals into empirically derived trajectories (7). In healthy females, LCMM identified three cortisol response types: mild, moderate, and heightened. Interestingly, both mild and heightened responders reported more negative affect than moderate responders (8). This finding addressed a long-standing challenge in the field, where traditional methods had faced difficulty linking cortisol and affective responses (9,10). In males, four cortisol profiles were identified—mild, moderately low, moderately high, and hyper-responses—but these did not differ in negative affect (11). In a mixed sample, LCMM identified two profiles: responders and non-responders with high baseline cortisol. Additional analyses showed that only responders successfully adjusted their behavior to changing reward contingencies in a probabilistic task (12). These findings illustrate that LCMM captures interindividual variability that may go beyond traditional metrics and reveal links to affect and behavior.

However, understanding of the processes that give rise to such variability is limited. A comprehensive review of rodent literature indicates that although cortisol is secreted peripherally, magnitude and duration of release is regulated centrally by brain regions including the amygdala, hippocampus, and medial prefrontal cortex (mPFC) (13). These regions are rich in MRs and GRs and participate in both the initiation and termination of the HPA axis through activation and inhibition of hypothalamic CRH neurons. Chronic stress disrupts this regulation, resulting in hyperactivity of the amygdala and impaired feedback in the hippocampus and mPFC, which were linked to exaggerated- and prolonged cortisol responses, respectively (13). Given the evolutionary conservation of the HPA system, these mechanisms likely apply to humans as well (13). However, most human studies have focused on aggregated cortisol metrics, rather than modeling trajectories over time.

Human studies linking brain structure and cortisol responses have mainly used structural and functional magnetic resonance imaging (sMRI, fMRI). sMRI research focuses on gray matter properties of brain regions, with subcortical volumes derived directly from anatomical segmentations and cortical thickness estimated via surface reconstructions to account for cortical folding (14). These measures have been associated with cortisol stress responses: smaller hippocampal volumes were related to both heightened and flattened cortisol stress responses (8,15). Amygdala volume correlated positively with cortisol stress responses in depressed adolescents, but negatively in healthy controls (16). Although cortical thickness has been studied less frequently in this context, reductions in mPFC thickness—especially medial orbitofrontal cortices (mOFC) and rostral anterior cingulate cortex (rACC)—are among the most robust findings in stress-related disorders (14,15).

fMRI studies typically compare neural activation under stress and control conditions. A prominent example is the Scan*STRESS* paradigm, which elicits psychosocial stress through mental rotation and arithmetic tasks performed under time pressure and social evaluation (16,17). A systematic review of such paradigms found that greater mPFC and ACC activation was associated with higher cortisol responses, while findings for the amygdala and hippocampus varied in direction (18). Some studies also reported time-dependent (exposure-time) effects in these regions, reflecting possible sensitization or habituation processes relevant to stress regulation (16,19–21). Collectively, these findings implicate the mPFC, hippocampus, and amygdala to play decisive roles in regulating cortisol release.

In summary, cortisol stress responses vary across individuals. While a rapid rise and recovery are generally considered adaptive, deviations from this pattern may confer latent vulnerability. LCMM approaches have captured meaningful subtypes of cortisol trajectories, linked to affect and behavior. However, the neurobiological mechanisms underlying these patterns remain insufficiently understood. Though rodent studies suggest a central HPA axis regulation via MR/GR-rich brain regions, most human studies lack the temporal specificity needed to link structure and function with cortisol dynamics. In the present study, we therefore investigated whether structural and functional parameters of key brain regions predict cortisol stress response trajectories identified through LCMM. We focused on four regions of interest (ROIs): the amygdala and hippocampus, based on their established relevance across species and imaging modalities (8,13,18,22,23), and two mPFC regions—the rACC and mOFC—which have been consistently implicated in structural (14,15) and functional approaches (18).

Following our pre-registration, for each ROI, we extracted subcortical volume or cortical thickness, task-based activation during Scan*STRESS*, and activation slopes reflecting exposure-time effects. We then used these parameters to predict cortisol trajectories defined by LCMM. We hypothesized that the inclusion of brain parameters may improve identification and prediction of cortisol trajectories beyond what is achieved in LCMMs without such parameters. In pre-registered exploratory analyses, we also investigated whether ROI-informed LCMMs identified negative affect trajectories, and whether these would overlap with cortisol trajectories. We hypothesized that interposition of brain parameters may help resolve the cortisol-affect incongruency observed using traditional methods. Finally, we explored psychometric properties of resulting trajectory groupings, as well as assignment coherence across models. These last analyses were not pre-registered but added to explore the functional meaning and reliability of identified trajectories.

## Materials and Methods

### Design

The study was pre-registered: https://osf.io/gzh2j/ and conducts a secondary analysis from five studies investigating healthy participants with Scan*STRESS* (*N* = 281; 145 females, mean age 25.21*±*7.65) (16,19,24–26). Scan*STRESS* was presented as a block-design with alternating stress and control blocks. Following the first half of the task (run1), participants received intercom feedback that their performance was below average, urging them to extend more effort to salvage data of the second half (run2) (see (16,17) for details). Studies followed a similar procedure, sampling cortisol and affect across ten timepoints surrounding stressor onset (*t* = −75, −15, −1, +15, +30, +50, +65, +80, +95, +110 *mins*), and acquiring T1-weighted structural images following the task (Figure 1A). Salivary cortisol samples were taken using salivette cotton swaps (Sarstedt, Nuembrecht, Germany). Affect was captured using the Positive and Negative Affect Schedule (PANAS) (27). We refer to the original publications for further details on data collection and our pre-registration for details on protocols and data selection.

**Figure 1.**
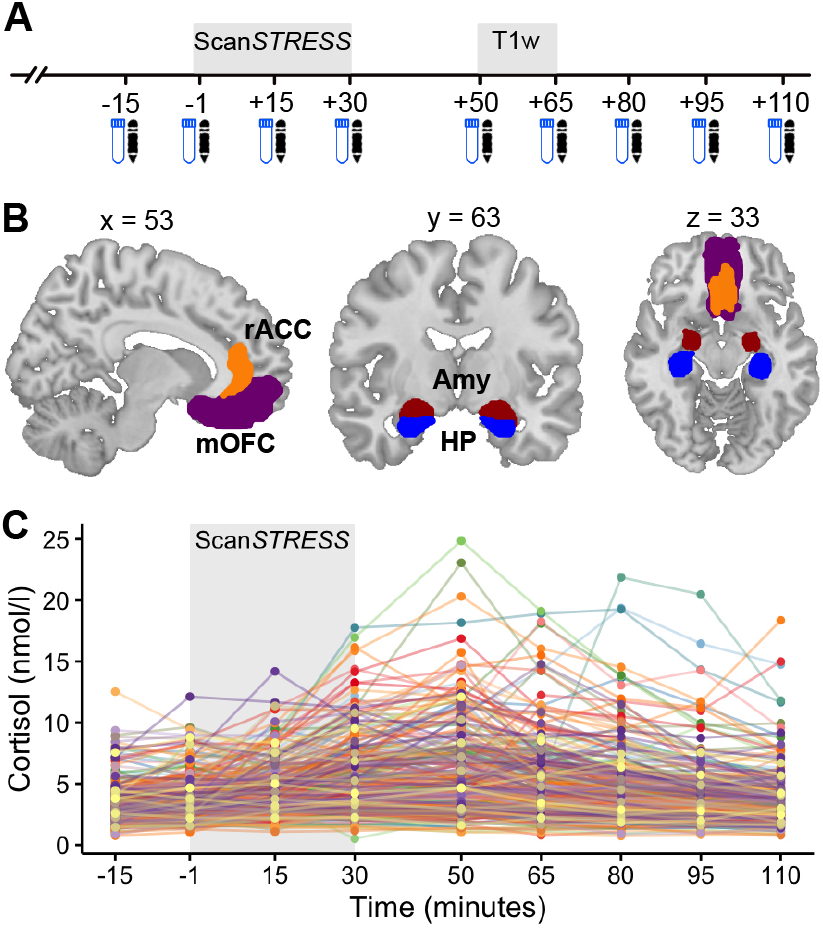
Experimental procedure, regions of interest (ROIs), and baseline variability prior to cortisol trajectory prediction. A) Experimental procedure consistent across studies. Blue icons represent salivary cortisol samples. Black pens represent negative affect samples. B) ROIs from which three parameters were extracted per participant (volume/thickness, activation, activation slope i.e., exposure-time effect). C) Variability in cortisol responses across the sample. Abbreviations: Amy, amygdala (red); HP, hippocampus (blue); mOFC, medial orbitofrontal cortex (purple); rACC, rostral anterior cingulate cortex (orange).

### (f)MRI preprocessing and analyses

All MRI data were acquired at Regensburg University, Germany, using a 3T Siemens MAGNETOM Prisma scanner with 64-channel head coil (Siemens Healthcare, Erlangen, Germany). Preprocessing and statistical analyses were performed using *HALFpipe* (28,29). Structural images were skull stripped using the convolutional neural network SynthSeg (30). Preprocessing involved motion correction (volume resampling), bias field correction, coregistration, registration to standard space, spatial smoothing (6mm), grand mean scaling, and high pass filtering at 125s. For statistical analyses, atlas-based parameters of structure, activation, and exposure-time effects were extracted in lateralized fashion for the four ROIs: amygdala, hippocampus, rACC, mOFC using Desikan-Killiany and Destrieux atlases (31,32). Structural parameters were derived from T1-weighted images via *FreeSurfer* surface and volume-based processing streams (i.e., volume for subcortical regions and thickness for cortical regions) (33,34). Functional parameters were derived from intraindividual contrasts within general linear models (*stress* > *control*). This was done in two ways: (1) globally over both runs of Scan*STRESS*, yielding mean activation, and (2) separately for each run, calculating the difference (*run*2 − *run*1), yielding slopes representing exposure-time effects. We note that we used motion regression rather than the pre-registered ICA-AROMA for model correction, as it performs less aggressive cleaning and potential data distortion (35). In summary, for each participant we derived three parameters per ROI: a structural parameter (volume (mm^3^)/thickness (mm)), mean activation (*z*-score), and exposure-time effect (*z*-score-slope). See Figure 1B for a visualization of ROIs.

### LCMM analyses

Figure 1C displays baseline variability in cortisol responses across the sample. To identify latent trajectories (i.e., groups of participants) exhibiting similar response trajectories, we fit latent process mixed models using the R package *lcmm* (v 2.1.0) (7,36). Model estimation was done using a grid of initial values to ensure convergence toward the global maximum (100 departures, 30 iterations) (7). First, we fit baseline models with 1-4 latent trajectories, including *age* and *sex* as covariates. This way, only residual heterogeneity in cortisol was explored after accounting for known factors of change over time (37,38). Note that we controlled for *sex* and not hormonal status as the latter had no influence on cortisol responses in our sample (see Supplementary section ‘hormonal status’).

In a second step, we explored whether adding brain parameters would improve model fit and predict cortisol trajectory assignment. To this end, 2-4 trajectory models were fit for each ROI, adding structure, activation, and exposure-time effect as predictors of trajectory membership (while still correcting for *age* and *sex*). Trajectory membership predictors entered a posterior polynomial logistic regression model, predicting cortisol trajectory from brain parameters. This required choice of a reference trajectory, against which the odds of belonging to each of the other trajectories were estimated (39).

For model comparison, we used common fit indices: Bayesian Information Criterion (BIC), size adjusted BIC (SABIC), and Akaike Information Criterion (AIC), which all indicate better fit with lower values. AIC and SABIC involve smaller penalties for model complexity. In samples < 400 individuals, BIC tends to penalize strongly for complexity in relation to sample size and is therefore biased towards selecting fewer trajectory models (40). Hence, the AIC is assumed to be the better selection criterion for our sample size (41). Entropy was consulted as a measure of discriminatory power, where values closer to one indicate clear separation into trajectories (41). Finally, models were also examined for biological plausibility and embedding with current theory (1–3).

To verify that trajectories of selected models were distinguishable, we ran mixed-effect analyses of variance (ANOVAs) with *time* as within- and *trajectory* as between-subjects factors. Though this approach may seem circular, it is considered the most appropriate post-hoc test for LCMMs (8,42). Note that we refer to trajectories as ‘hyper’ and ‘hypo’ responses to reflect relative sample differences and maintain consistent terminology across studies (8,11), but do not imply clinical (mal)adaptation.

In non-pre-registered analyses, we further explored the functional meaning of identified trajectories using Chi-square tests and mixed-effects ANOVAs on *age, sex*, trait-like affect using Beck Depression Inventory (*BDI-II*) (43) and Anxiety Sensitivity Index (*ASI*) (44), as well as indices of chronic stress using the screening scale of the Trier Inventory for the assessment of Chronic Stress (*TICS*) (45), Life Events Checklist (*LEC*) (46), and Childhood Trauma Questionnaire (*CTQ*) (47). Using Cohen’s and Fleiss’ kappa, we also explored trajectory assignment coherence across different models (i.e. between ROIs and cortisol/affect).

Due to convergence issues, we deviated from our pre-registration in two additional points. First, we omitted total brain volume correction in subcortical models as this would have corrected cortisol trajectories, rather than brain parameters, and caused convergence issues, likely due to collinearity with *sex*. Secondly, we had planned to estimate separate models for left and right ROIs. However, this complicated model comparison as lower-entropy solutions showed limited assignment coherence across sides, despite visually comparable trajectories. To circumvent this, we first included a side variable as interaction terms of brain parameters, which increased complexity and non-convergence. Finally, given high correlations between left and right parameters (Supplementary Figure S1), we merged sides per ROI, aiding convergence and model selection. Following model estimation, we nevertheless plotted left and right parameters individually, yielding similar patterns across sides (see Figures 2, 3, and 5 C-E). The R code of *lcmm* analyses is provided here: https://osf.io/gk6w9/.

**Figure 2.**
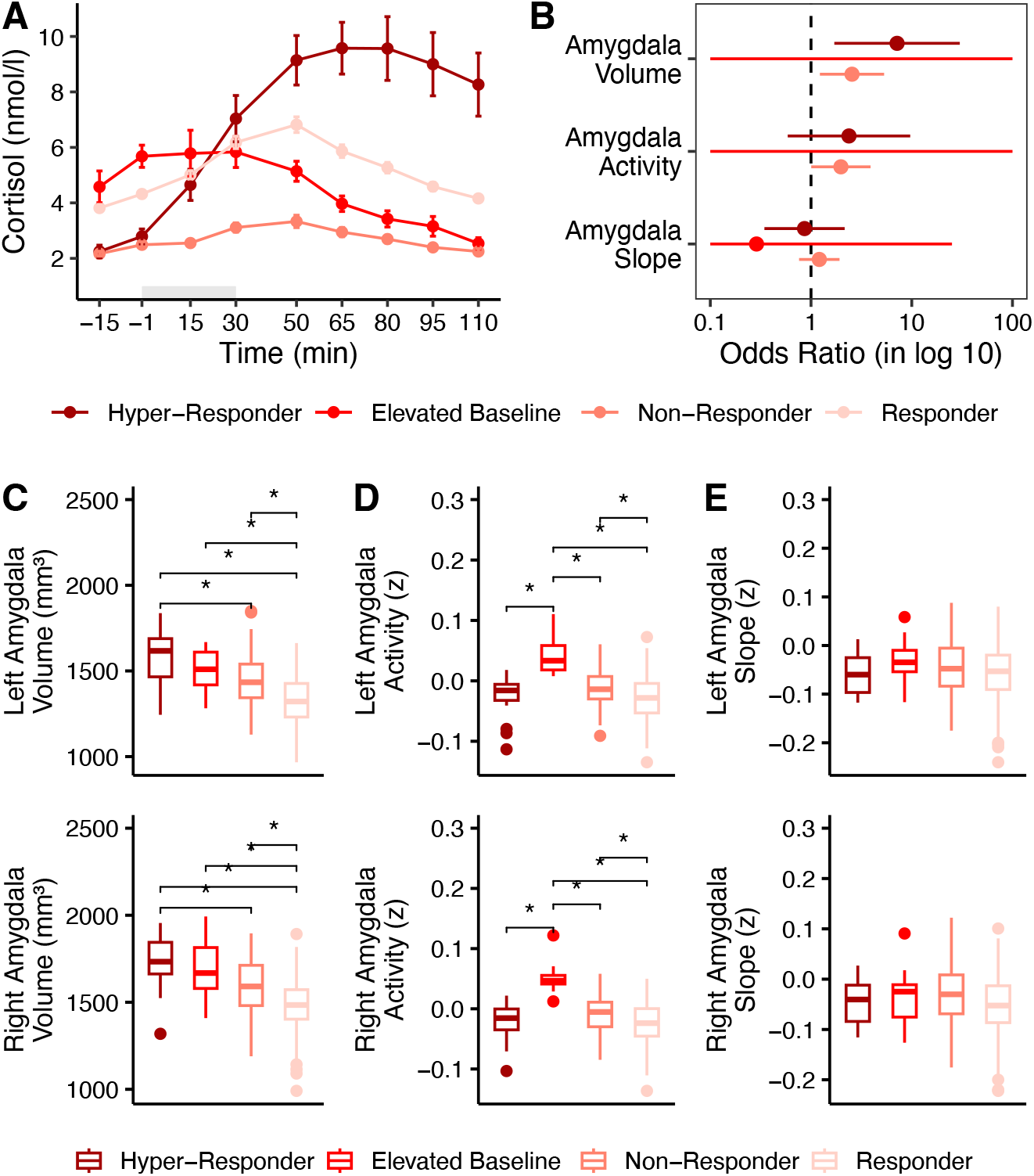
A) Mean cortisol of the four-trajectory amygdala model. Grey shading indicates Scan*STRESS* timing. B) Amygdala parameters as posterior predictors of trajectory membership. Odds Ratios (ORs) were computed as compared to the reference trajectory (‘Responder’). The analysis accounted for the uncertainty (entropy) in trajectory assignment. Note that confidence intervals of the ‘Elevated Baseline’ trajectory were large and clipped at [0.1, 100] for plotting. C-E). Boxplots of amygdala parameters (volume, activity, slope (i.e., exposure-time effect), respectively) across trajectories. For each plot, left amygdala parameters were displayed on top, right parameters below. Within plots, statistically significant pairwise comparisons are displayed while non-significant comparisons were left blank.

**Figure 3.**
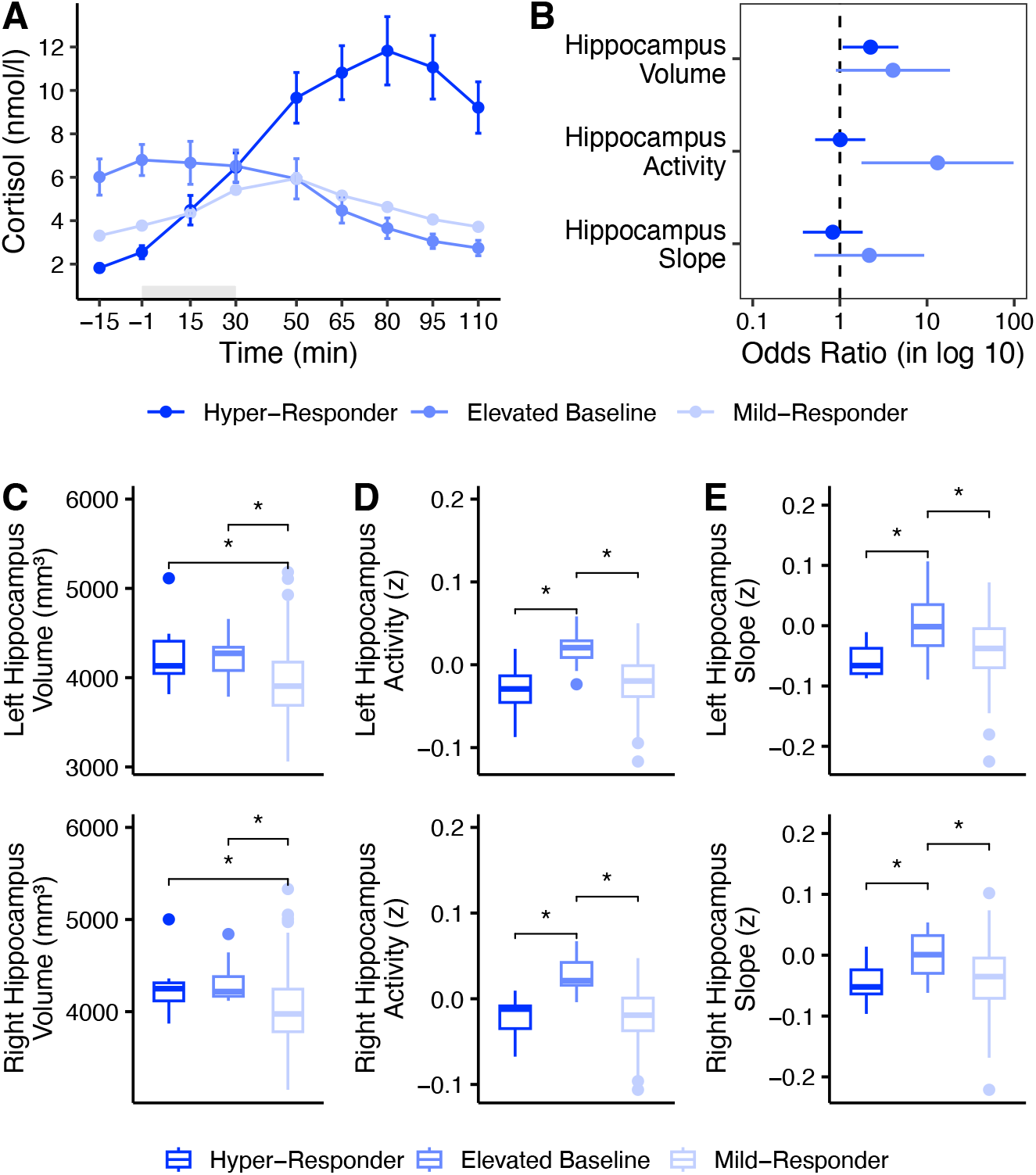
A) Mean cortisol of the three-trajectory hippocampus model. Grey shading indicates Scan*STRESS* timing. B) Hippocampus parameters as posterior trajectory membership predictors. Odds Ratios (ORs) were computed as compared to the reference trajectory (‘Mild Responder’). The analysis accounted for the uncertainty (entropy) in trajectory assignment. C-E) Boxplots of hippocampus parameters (volume, activity, slope (i.e., exposure-time effect), respectively) across trajectories. For each plot, left hippocampus parameters were displayed on top, right parameters below. Within plots, statistically significant pairwise comparisons are displayed while non-significant comparisons were left blank.

### Exploratory affect analyses

As pre-registered exploratory affect analyses, we first entered cortisol trajectory groupings from selected brain models into mixed-effect ANOVAs on PANAS negative affect, with *time* as within- and *trajectory* as between-subjects factors. When cortisol trajectory groupings did not explain variance in negative affect, we tested whether re-running LCMMs on negative affect would identify affect trajectories that corresponded meaningfully with cortisol trajectories. Therefore, we replicated baseline and brain models described in the ‘LCMM analyses’ section using negative affect as the outcome.

## Results

### Baseline models

When comparing the four baseline cortisol models (corrected for *sex* and *age*), all model comparison indices (AIC, BIC, SABIC, Entropy) favored the single-trajectory solution (Table 1). In the following, we will compare models with brain parameters to this best-performing baseline model.

**Table 1.**
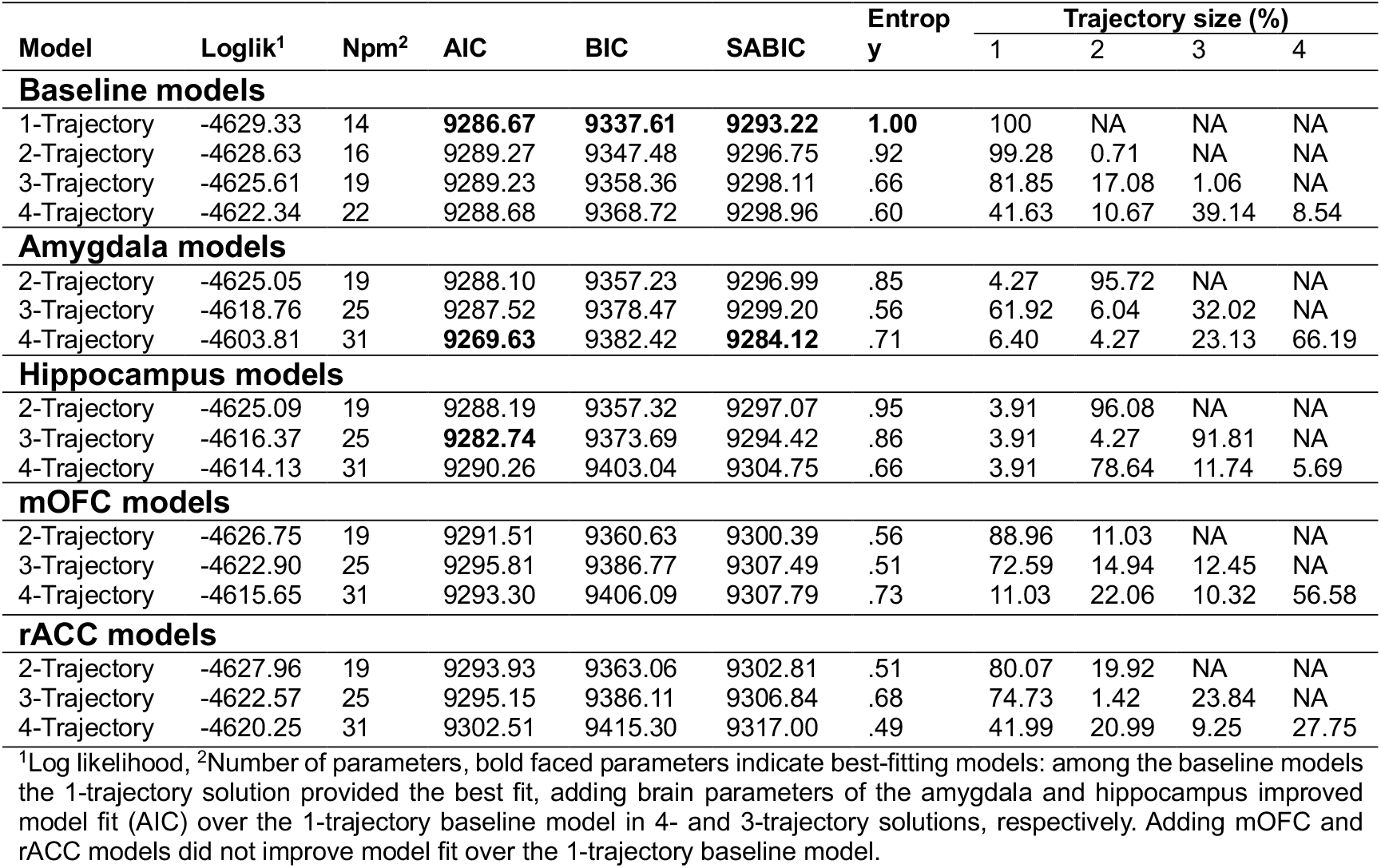
Between model comparison of cortisol models.

### Brain models

Among amygdala models, AIC and SABIC favored the four-trajectory model (*entropy* =.71, Table 1). The model aligned well with current theory (3,4) and showed similarity with existing literature on latent cortisol trajectories (8,11). See Figure 2A for a depiction of the selected model. A mixed-effect ANOVA verified that the cortisol trajectories were distinguishable (main effect of time: *F*(3.20, 822.94) = 26.70, *p* <.00, 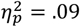, main effect of trajectory: *F*(3.00, 257.00) = 37.85, *p* <.00, 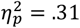, time*trajectory interaction *F*(9.61, 822.94) = 15.26, *p* <.00, 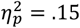). The first trajectory showed a heightened cortisol stress response with slowed recovery (‘hyper-responder’, *N* = 18), while the second trajectory was characterized by higher basal cortisol levels and no stress response (‘elevated baseline’, *N* = 12). The third trajectory exhibited a non-response profile (‘non-responder’, *N* = 65) while the fourth trajectory showed a moderate cortisol stress response, with recovery to baseline two hours later (‘responder’, *N* = 184). For posterior trajectory membership prediction, we selected this responder trajectory as the reference, as it represents adaptive response (4), to which aberrant responses could be compared. Results showed that higher amygdala volume increased the odds of a ‘hyper-response’ (*coef* = 1.96, *SE* =.73, *Wald* = 2.68, *p* <.01, *OR*[95%*OR*−*CI*] = 7.14[1.70, 29.97]) or ‘non-response’ (*coef* = 0.93, *SE* =.37, *Wald* = 2.50, *p* =.01, *OR*[95%*OR*−*CI*] = 2.55[1.22, 5.31]). Amygdala activation also predicted trajectory membership: Compared to ‘responders’, higher amygdala activation increased the odds of a cortisol ‘non-response’ (*coef* =.68, *SE* =.34, *Wald* = 1.97, *p* =.05, *OR*[95%*OR*−*CI*] = 1.97[1.01 3.89]). See Figure 2B for a depiction of amygdala parameter ORs per trajectory as well as Figures 2C-E and Supplementary Table S1 for mean amygdala parameters stratified by side and trajectory). Supplementary Table S2 shows that ‘hyper-responder’ and ‘elevated baseline’ groups were predominantly male, while no group differences were observed on other questionnaires.

Concerning the hippocampus models, AIC favored the three-trajectory model with good entropy of. 86 (Table 1) and theoretical plausibility (3,4) (Figure 3A). A mixed-effect ANOVA again verified that trajectories were distinguishable (main effect of time: *F*(3.11, 802.97) = 22.91, *p* <.00, 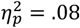, main effect of trajectory: *F*(2.00, 258.00) = 12.23, *p* <.00, 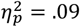, time*trajectory interaction: *F*(6.23, 802.97) = 28.92, *p* <.00, 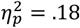). The first trajectory showed a heightened cortisol response with slowed recovery (‘hyper-responder’, *N* = 11) and the second trajectory exhibited elevated baseline cortisol but non-response to the stressor (‘elevated baseline’, *N* = 12). The third trajectory showed a mild cortisol stress response, with rapid return to baseline two hours later (‘mild responder’, *N* = 254). We again chose the ‘mild responder’ trajectory as the reference for posterior polynomial logistic regression. Results showed that higher hippocampus volume was related to higher odds of cortisol ‘hyper-response’ (*coef* =.80, *SE* =.37, *Wald* = 2.14, *p* =.03, *OR*[95%*OR*−*CI*] = 2.24[1.07, 4.71]). Additionally, higher hippocampus activation increased the odds of membership in the ‘elevated baseline’ over the ‘mild-responder’ trajectory (*coef* = 2.57, *SE* = 1.02, *Wald* = 2.51, *p* =.01, *OR*[95%*OR*−*CI*] = 13.18[1.76, 98.33]). See Figure 3B for a depiction of hippocampus parameter ORs per trajectory. Figures 3C-E and Supplementary Table S3 show hippocampus parameter means across sides and trajectories. Table 2 shows that ‘hyper-responder’ and ‘elevated baseline’ groups were predominantly male. Additionally, the ‘elevated baseline’ group reported significantly higher *CTQ* scores than other trajectories, though still remaining in the ‘low to moderate’ severity quintile (48). No differences were observed on other variables.

**Table 2.** Characterization of hippocampus model derived cortisol classes.

|  | Trajectory mean (SD) |  |  | Statistic(df) | p | Effect size |
| --- | --- | --- | --- | --- | --- | --- |
|  | Hyper-responder | Elevated baseline | Mild-responder |  |  |  |
| Age | 23.45 (±3.11) | 28.50 (±9.01) | 25.14 (±7.70) | $F(2, 278) = 1.41$ | .24 | $\eta^2 = .01$ |
| Sex | | | | $\chi^2(2) = 16.34$ | < .01* | $V = .24$ |
| M | 11 | 9 | 116 |  |  |  |
| F | 0 | 3 | 142 |  |  |  |
| BDI-II | 9.18 (±10.03) | 11.25 (±9.62) | 13.56 (±11.69) | $F(2, 272) = 0.95$ | .38 | $\eta^2 = .01$ |
| TICS | 15.64 (±6.28) | 13.50 (±7.62) | 18.00 (±9.63) | $F(2, 276) = 1.57$ | .20 | $\eta^2 = .01$ |
| ASI (Total) | 15.00 (±7.54) | 19.67 (±11.44) | 20.13 (±10.92) | $F(2, 273) = 1.19$ | .30 | $\eta^2 = .01$ |
| CTQ (Total) | 29.00 (±4.34) | 37.00 (±10.55) | 32.15 (±7.35) | $F(2, 274) = 3.54$ | .03*# | $\eta^2 = .03$ |
| LEC (Total) | 67.36 (±6.02) | 70.75 (±8.07) | 67.09 (±12.74) | $F(2, 278) = 0.50$ | .60 | $\eta^2 = .00$ |

Abbreviations: ASI = Anxiety Sensitivity Index, BDI-II = Beck Depression Inventory II, CTQ = Childhood Trauma Questionnaire, LEC = Life Events Checklist, TICS = Trier Inventory for the Assessment of Chronic Stress (Screening Scale).
\* indicate statistically significant differences between groups. # Pairwise comparisons: hyper-responder < elevated baseline ( $p = .02$ ), hyper-responder < mild-responder ( $p = .04$ ).

For the mOFC and rACC models, none of the fit indices favored higher trajectory numbers over the single-trajectory solution without additional brain predictors (Table 1). We therefore decided not to explore these models further.

Finally, we assessed the overlap between amygdala and hippocampus models. Visual inspection (Figures 2A and 3A) suggested that the amygdala ‘responder’ and ‘non-responder’ trajectories had merged into one hippocampus ‘mild responder’ trajectory. To test this, we merged amygdala ‘responders’ and ‘non-responders’ and compared this to hippocampus assignments, yielding moderate agreement (*Cohen*’*s Kappa* =.57, *z* = 12.50, *p* <.00). This indicates substantial overlap between trajectories across models, with only a few individuals reclassified (Figure 4).

**Figure 4.**
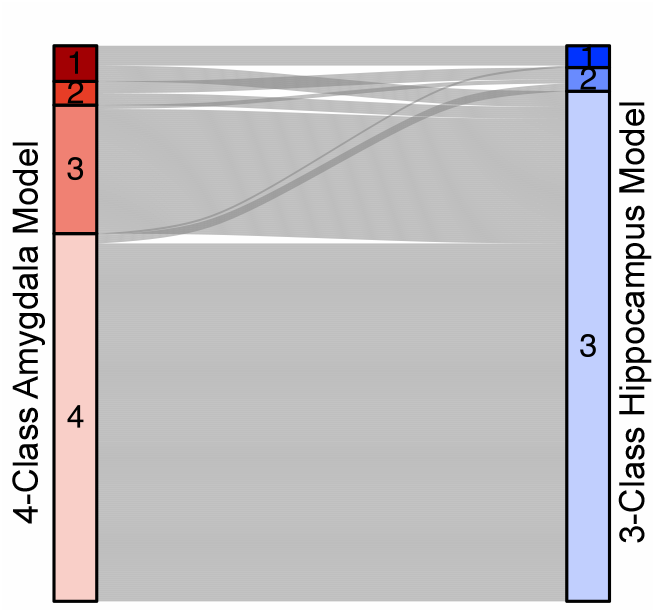
Alluvial plot of participant flow between amygdala (left) and hippocampus (right) cortisol models.

### Exploratory affect analyses

Next, we explored whether cortisol groups derived from amygdala and hippocampus brain models would explain variance in stress-related negative affect. Mixed-effects ANOVAs indicated that brain-derived cortisol trajectories did not differ in negative affect (all *p*’*s* >.05). We then explored whether running baseline and brain models on negative affect data would identify affect trajectories that corresponded with cortisol trajectories (see Supplementary section ‘exploratory affect analyses’ for statistical detail). Results indicated that while distinct negative affect trajectories were identified with and without brain predictors, these did not overlap with cortisol trajectories (Supplementary Table S4, Figure S3, Figure S4). Additionally, amygdala, hippocampus, and rACC predictors showed no utility in predicting negative affect trajectories (all *p*’*s* >.05). However, among mOFC affect models, AIC and SABIC favored the four-trajectory model with *entropy* =.67 (Supplementary Table S3, Figure 5A). The four trajectories followed a similar pattern but varied in negative affect intensity: ‘strong-lasting’ (*N* = 21), ‘moderate-lasting’ (*N* = 69), ‘moderate-brief’ (*N* = 69), and ‘mild-brief’ (*N* = 109). For posterior polynomial logistic regression, we chose the ‘mild-brief’ trajectory as the reference due its resemblance to ‘positive appraisal styles’ associated with resilience (10). Results showed that compared to this reference, greater mOFC thickness increased the odds of ‘moderate-lasting’ trajectory assignment (*coef* =.64, *SE* =.29, *Wald* = 2.17, *p* =.02, *OR*[95%*OR*−*CI*] = 1.90[1.06, 3.40]). See Figure 5B for mOFC odds ratios and Figures 5C-E for descriptive boxplots stratified by side and trajectory. Questionnaire analyses indicated that the intensity of acute stress-related affect was closely mirrored by trait-like stress propensity: while the ‘mild-brief’ trajectory scored lowest on depression, anxiety, life stress, and childhood trauma, more pronounced negative affect trajectories had incrementally higher scores on these scales (Table 3). We do note that this model had an entropy of. 67, indicative of trajectory assignment ambiguity. Though ORs displayed in Figure 4B are corrected for such ambiguity, we nevertheless urge a cautious interpretation of these results.

**Figure 5.**
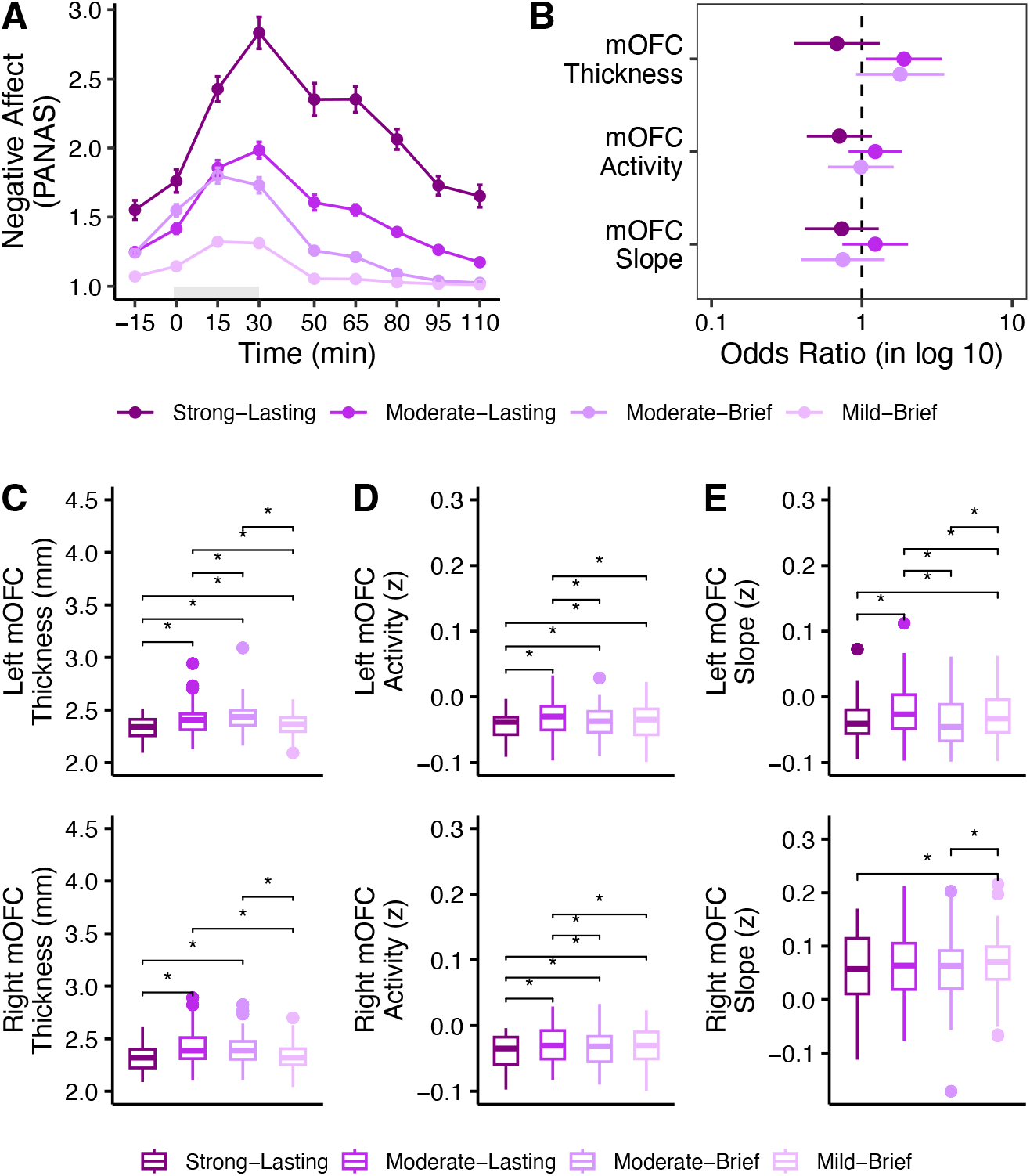
A) Medial orbitofrontal cortex (mOFC)-derived negative affect trajectories. Grey shading indicates Scan*STRESS* timing. B) mOFC parameters (thickness, activity, slope (i.e., exposure-time effect)) as posterior trajectory membership predictors. Odds Ratios (ORs) were computed as compared to the healthy reference trajectory (‘Mild-Brief’). The analysis accounted for the uncertainty (entropy) in trajectory assignment. C-E) Boxplots of mOFC parameters across trajectories. For each plot, left mOFC parameters were displayed on top, right parameters below. Statistically significant pairwise comparisons are displayed within plots, non-significant ones were left blank.

**Table 3.**
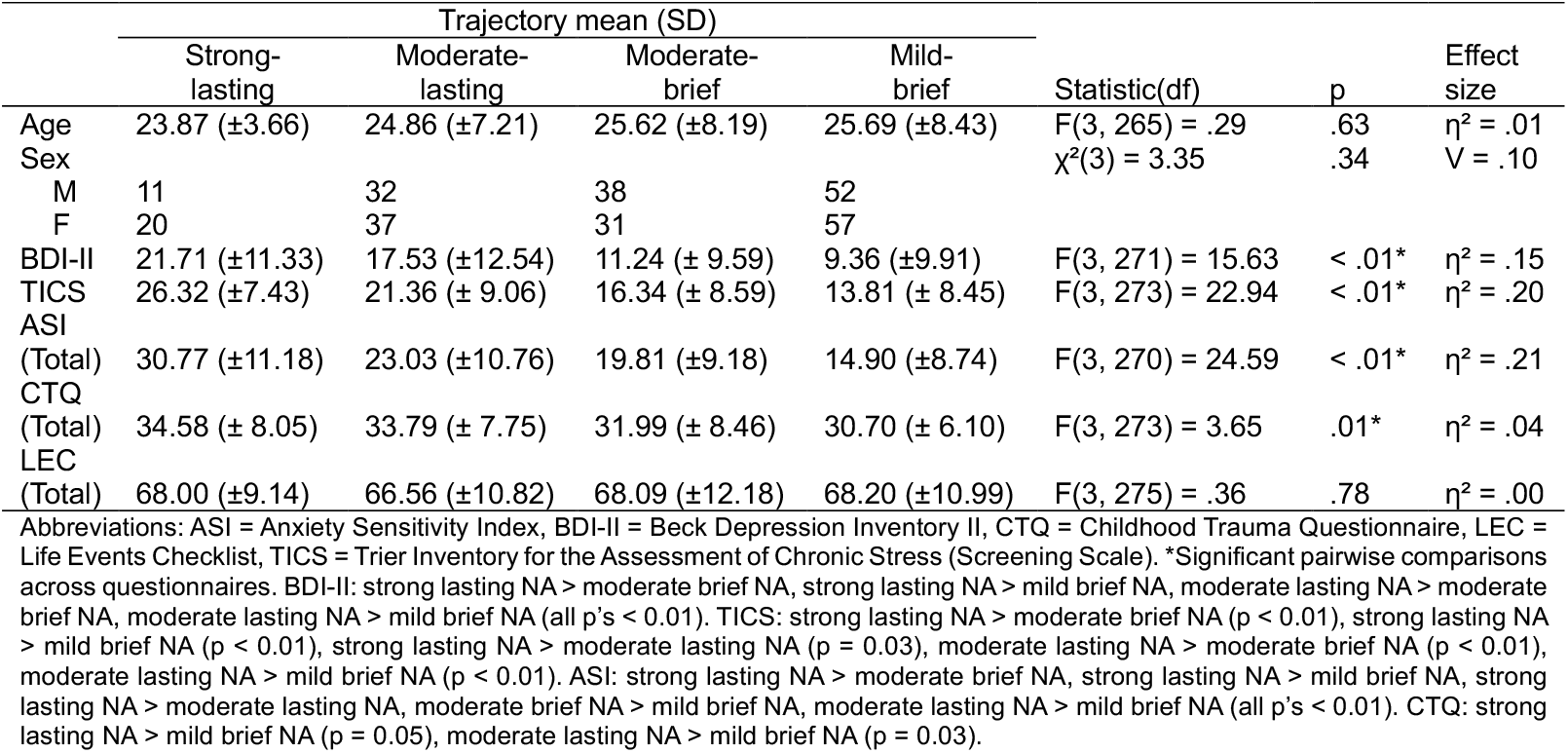
Characterization of mOFC model-derived negative affect trajectories.

## Discussion

Varied cortisol stress responses can be captured using LCMM. Here, we explored whether adding functional and structural brain parameters from the amygdala, hippocampus, rACC, and mOFC would aid the identification of cortisol stress response trajectories. Without brain predictors, the single-trajectory model showed the best fit, indicating largely homogeneous cortisol responses in the sample. The inclusion of amygdala and hippocampus parameters improved model fit and yielded more nuanced solutions (three and four trajectories, respectively), suggesting that interindividual differences in limbic structure and function meaningfully contributed to the identification of divergent stress responses. In contrast, models incorporating rACC and mOFC parameters did not outperform the baseline cortisol model. However, mOFC parameters did aid the identification of affect trajectories with similar pattern but varying intensity. We elaborate on these findings in the sections below.

### Amygdala parameters and cortisol trajectories

The best-fitting amygdala model identified four cortisol response trajectories, which were comparable to previous investigations in number and size (8,11). Greater amygdala volume increased the odds of a cortisol ‘hyper-response’, which aligns with its key role in HPA-axis activation via disinhibition of hypothalamic CRH-neurons (13). In rodents, prolonged stress leads to dendritic growth and increased spine density in the amygdala, promoting vigilance and stress responsivity (49). In humans, this may manifest as greater amygdala volume observed during early disease development, such as following greater life stress (50), in subclinical trait anxiety (51), or in adolescent depression (23). Other studies also reported exaggerated cortisol stress responses following prolonged life stress (11) and in depressed adolescents with larger amygdala volumes (23). Together, these results may suggest that enlarged amygdala volumes and exaggerated cortisol release reflect early signs of stress sensitization. However, hyper-response individuals in our sample did not report greater life stress, psychiatric symptoms, or negative affect than other trajectories, possibly reflecting disturbed stress systems before subjective symptoms emerge. Given that cortisol hyper-responses prospectively predict adverse health outcomes (3,4), longitudinal studies should further disambiguate the developmental significance of enlarged amygdala volumes in relation to cortisol stress responses.

Interestingly, higher amygdala volume and activation also increased the odds of cortisol ‘non-responses’. One possible interpretation is that this pattern reflects a latent vulnerability. Amygdala hypertrophy has been linked to stress sensitization in both rodents and humans (49,50) and amygdala hyperactivity prospectively predicts PTSD symptoms (52,53). PTSD is also commonly associated with flattened cortisol stress responses, likely due to feedback hypersensitivity mitigating neurotoxic effects of prolonged cortisol exposure (3). In line with this, young adults with a history of early life adversity have shown larger amygdala volumes along with blunted cortisol stress responses (54). In this context, our findings may indicate amygdala hypertrophy and hyperactivity initiating the HPA axis, while its output is dampened at the adrenal level. Alternatively, this trajectory might also reflect healthy adaptation. As a central hub of the salience network, the amygdala is typically activated during stress, facilitating attentional reorientation and selection of habitual coping strategies (2,21,55). Cortisol non-responses associated with larger volumes and higher activity could therefore also reflect a healthy response to a mild stressor, with a relative preservation of neural integrity due to lesser cortisol exposure. In line with this, healthy adolescents with larger amygdala volumes also exhibited lower stress-related cortisol (23).

In summary, greater amygdala volume predicted cortisol hyper-responses, suggesting that amygdala hypertrophy and exaggerated cortisol reactivity may serve as early markers of stress sensitization, emerging even before subjective symptoms appear. At the same time, greater amygdala volume and activation also predicted cortisol non-responses. This pattern may indicate either an enlarged and hyperactive amygdala driving HPA axis activity to a point of adrenal exhaustion, or, conversely, a relatively intact amygdala maintained by a mild cortisol stress response propensity and reduced neurotoxic exposure.

### Hippocampus parameters and cortisol trajectories

The hippocampus-based model identified three cortisol response trajectories which largely overlapped with those derived from the amygdala model. Higher hippocampal volume also increased the odds of cortisol ‘hyper-responses’, which aligns with the hippocampus’ role in GR-mediated feedback. Following prolonged stress, feedback sensitivity may diminish and result in prolonged cortisol release observed in anxiety and depression (3). However, the relationship between these conditions and hippocampal volume is opposing: while depression is often associated with reduced volume (14,56), anxiety disorders tend to be accompanied by larger volumes (51,57,58). Based on this, some authors have speculated that stress-related limbic plasticity may be biphasic along disease development: with early hypertrophy followed by later atrophy (51). Given that individuals in the hyper-response trajectory were otherwise healthy, volume increases may reflect such early-stage hypertrophy and feedback insensitivity. This would align with findings linking larger hippocampal volumes to subclinical anxiety (51), which is characterized by enhanced contextual fear-learning (57). However, previous studies have also found smaller hippocampal volumes in relation to heightened cortisol stress responses in healthy individuals, which was interpreted as neurotoxic damage following excessive cortisol exposure (8,22). Notably, neither those studies nor ours found greater self-reported life stress in hyper-responders. In contrast, we found them to report the lowest levels of childhood trauma, which could suggest that this trajectory reflects a healthy phenotype with efficient stress system mobilization, rather than the dysregulation suggested above.

Finally, higher hippocampal activity predicted membership in the ‘elevated baseline’ trajectory. Unlike the amygdala, the hippocampus is typically deactivated during stress (2,55), possibly reflecting an increased signal-to-noise ratio where only a few cells are active to encode the experience with high specificity (59). Conversely, less selective hippocampal cell activation may lead to overgeneralized memories (59), such as those seen in PTSD (60) and anxiety (61). Indeed, trauma-exposed individuals displaying sustained threat-related hippocampal activity later developed more PTSD symptoms (62). Together with flattened cortisol stress responses being associated with feedback hypersensitivity (3), individuals in this trajectory may be exposed to a ‘double hit’ of overgeneralized memory encoding and impaired HPA axis upregulation. Supporting this interpretation, the ‘elevated baseline’ trajectory reported higher levels of childhood trauma, consistent with models proposing that cortisol non-responsivity emerges secondarily to prolonged (early) stress exposure (3,63).

In summary, larger hippocampal volumes were associated with cortisol hyper-responsivity, which may either reflect hypertrophy and stress sensitization characteristic of early disease stages, or a preserved and flexible stress system in individuals without early traumatic experiences. By contrast, increased hippocampal activation predicted elevated baseline non-response trajectories, possibly reflecting maladaptive encoding and feedback hypersensitivity following early adversity. These findings are broadly consistent with biphasic models of disease development, in which early increases in limbic volume and cortisol reactivity are followed by later limbic atrophy and attenuated stress reactivity.

### Exploratory affect analyses

In our exploratory affect analyses, neither amygdala-nor hippocampus-derived cortisol trajectories differed in stress-induced negative affect. Similarly, ROI-derived negative affect trajectories did not mirror cortisol-based response trajectories. These findings align with a body of literature reporting inconsistent associations between cortisol and affective stress responses (9,11). In fact, the two systems may even be functionally dissociated, as dexamethasone administration prior to psychosocial stress suppressed cortisol reactivity but left affective responses intact (64,65).

In ROI-derived negative affect models, amygdala, hippocampus, and rACC parameters did not predict trajectories, but mOFC parameters did. The mOFC model had four trajectories ranging from ‘mild-brief’ to ‘strong-lasting’ negative affect, which paralleled participants’ trait-like affect. Greater mOFC thickness increased the likelihood of ‘moderate-lasting’ negative affect, aligning with the mOFC’s role in emotion regulation via inhibitory projections to the amygdala (66). Previous studies have linked greater mOFC thickness to reduced trait anxiety following childhood adversity (67) as well as resilience in adolescents exposed to peer abuse (68). In our sample, the ‘moderate-lasting’ trajectory reported higher levels of childhood trauma, while their psychiatric symptoms remained in the mild-to-moderate range (43,44). This aligns with definitions of resilience as mental health despite adversity (69) and with stress inoculation theories positing that moderate levels of early adversity may foster later resilience (70). Indeed, monkeys exposed to controlled stressors in early life exhibited enhanced adult stress resilience, which was accompanied by greater mOFC volume and stronger mOFC–amygdala connectivity (71,72). Together, these results suggest that greater mOFC integrity fosters acute affect regulation, which in turn may support resilience over time.

In sum, our results further corroborate a dissociation between cortisol and affective responses to stress. While cortisol trajectories were best explained by adding limbic parameters, affective trajectories were predicted by mOFC structure. Particularly, higher mOFC thickness was linked to moderate yet lasting negative affect in a potentially resilient group. These findings align with previous work positioning the mOFC as a key site for affect regulation and resilience.

### Limitations and future direction

Despite the considerable strengths of our study, including a large sample size and data-driven LCMM-approach to investigate cortisol variability alongside limbic and mPFC (s/f)MRI parameters, we were unable to control for three potentially relevant variables. First, though we included *sex* as a covariate, which accounted for systematic sex differences, hyper-response trajectories nevertheless consisted exclusively of males, which generally release more stress-related cortisol (37). To circumvent such conflations, studies in larger samples may aim to investigate sexes separately or include hormonal status as a covariate (73).

Secondly, we omitted total brain volume as a separate covariate, as the collinearity between sex and brain size appeared to cause non-convergence. However, as brain size influences the size of subcortical regions (74) this may have influenced our results.

Thirdly, though we had planned to investigate lateralization effects, convergence issues led us to merge ROIs across hemispheres. However, as lateralization effects were reported in both functional (75) and structural (76) investigations of acute stress, larger studies should include interaction terms for hemispheres.

Furthermore, exposure-time effects did not predict cortisol or affect trajectories. Changes over the course of stress exposure may thus not explain additional variance over mean activation in healthy samples. However, since such neural sensitization and habituation effects were shown in clinical samples (16,19–21), this parameter may prove relevant in future clinical investigations.

Moreover, instead of estimating separate cortisol and affect models, investigations combining both outcomes found it was exactly the mismatch (flattened cortisol with heightened negative affect) which was informative of clinical status (77). Exploiting integrational methods such as multlcmm (39) may therefore pose an exciting future avenue to advance understanding of cortisol-affect dissociations.

Finally, utilizing the standard questionnaire battery implemented with Scan*STRESS* may not have been most appropriate to characterize trajectories. Especially the *CTQ* has been critiqued for its incompleteness regarding range and timing of exposure (78), and conflation with recall bias (79). To better explore associations between childhood adversity and cortisol trajectories, future investigations should characterize adversity using more comprehensive instruments (80).

## Conclusion

The identification and prediction of stress response trajectories was improved by adding brain parameters from the amygdala, hippocampus, and mOFC. While limbic parameters informed cortisol responses, mOFC parameters informed both acute and trait-like negative affect responses. Though these findings provide first insights into how MRI-derived parameters of HPA-regulating regions may relate to aberrant cortisol and affect responses, the directionality and functional meaning of these results awaits disambiguation in longitudinal designs.

## Contributors

CRediT: conceptualization G-IH, RL, methodology G-IH, RL, LK, formal analysis RL, G-IH, LK, investigation G-IH, CB, MG, JK, HLP, NS, data curation G-IH, LW, writing - original draft RL, G-IH, writing - review & editing G-IH, LK, CB, MG, JK, HLP, NS, LW, BMK, SW, HW, visualization RL, supervision G-IH, funding acquisition: G-IH.

## Data availability statement

The datasets generated and/or analyzed during the current study are available from the corresponding author on reasonable request. A repository of studies that have already used and published the data is available here: https://osf.io/echja/.

