## Supplementary Material for "Predicting stress response trajectories: Differential contributions of limbic and prefrontal regions to cortisol and affective responses"

**Abstract**

Why do individuals respond differently to stress? Since rodent studies indicated that stress regulation relies on limbic and medial prefrontal cortex (mPFC) outputs, we aimed to investigate whether data from these regions could also predict cortisol and affect trajectories following psychosocial stress in humans. In this pre-registered study, 281 healthy adults (145 female) were exposed to Scan*STRESS*. Repeated assessments of salivary cortisol and negative affect were used to identify response trajectories (i.e. groups of participants) using latent class mixture modelling (LCMM). LCMMs without brain predictors were compared to LCMMs including structural (volume, thickness) and functional (activation, exposure-time effect) predictors from the amygdala, hippocampus, or mPFC regions. Results showed that cortisol LCMMs without brain predictors exhibited a single mean trajectory. Adding brain predictors resulted in three to four response trajectories, depending on region and outcome. Within identified models, cortisol 'hyper-response' trajectories were predicted by larger amygdala and hippocampus volumes. Cortisol 'non-responses' were predicted by greater amygdala activation and volume. 'Elevated baseline' cortisol was predicted by higher hippocampal activation. mPFC markers did not predict cortisol trajectories, however, medial orbitofrontal cortex parameters identified affect response profiles mirroring trait-like affect. Together, our findings suggest dissociated roles of limbic and mPFC regions in stress regulation: While limbic structures predicted cortisol responses, the mPFC shaped affective experience.

**Brain model supplements**

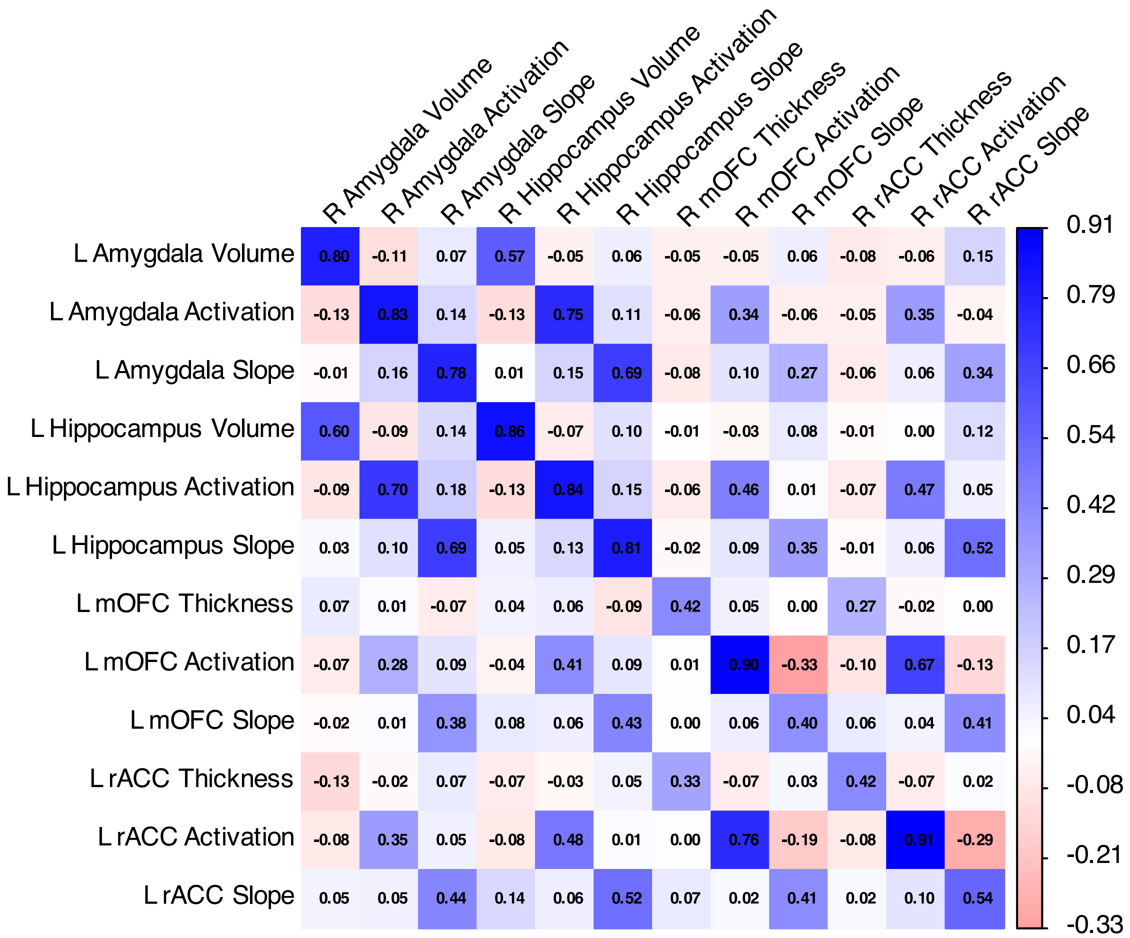

**Figure S1.** Correlation matrix of left and right parameter estimates of each region of interest (ROI). Note that the slope parameter refers to the ‘exposure-time effect’. Abbreviations: L, left; R, right; mOFC, medial orbitofrontal cortex; rACC, rostral anterior cingulate.

*Hormonal status*

Our brain models were controlled for sex and not hormonal status (i.e., cycle phase or contraceptive use) even though prior work showed cortisol differences as a function of these variables (1). This is because mixed effects analyses of variance (ANOVAs) in our sample indicated that while there were differences in cortisol between sexes (main effect of time: $F(3.08, 797.33) = 67.33, \eta_p^2 = .20$, main effect of sex: $F(1, 259) = 18.31, p < 0.01, \eta_p^2 = .06$, sex*time interaction: $F(3.08, 797.33) = 17.02, p < .01, \eta_p^2 = .06$), no differences were observed between cycle phases of females (main effect of time: $F(3.02, 431.77) = 16.79, p < .01, \eta_p^2 = .11$), main effect of cycle: $F(1, 143) = .15, p = .69, \eta_p^2 = .00$, cycle*time interaction: $F(3.02, 431.77) = 1.70, p = .16, \eta_p^2 = .01$).

**Table S1.** Mean amygdala predictor values across amygdala-derived cortisol trajectories

| | Trajectory mean (SD) | | | | | | |
|---|---|---|---|---|---|---|---|
| | Hyper-responder | Elevated baseline | Non-responder | Responder | Statistic(df) | *p* | *η²* |
| Volume (mm³) | | | | | | | |
| L | 1588.23 (179.31) | 1507.38 (122.97) | 1455.07 (157.41) | 1332.69 (138.93) | F(1, 279) = 79.55 | < .01* | .22 |
| R | 1734.60 (159.73) | 1691.49 (177.87) | 1595.58 (154.57) | 1485.72 (150.18) | F(1, 279) = 71.24 | < .01* | .20 |
| Activity (z) | | | | | | | |
| L | -.03 (.03) | .04 (.03) | -.01 (.03) | -.03 (.04) | F(1, 279) = 10.89 | < .01* | .04 |
| R | -.02 (.03) | .05 (.03) | -.01 (.03) | -.02 (.03) | F(1, 279) = 17.02 | < .01* | .06 |
| Slope (z) | | | | | | | |
| L | -.06 (.04) | -.03 (.05) | -.04 (.06) | -.05 (.06) | F(1, 279) = .47 | .49 | .00 |
| R | -.05 (.04) | -.04 (.06) | -.03 (.06) | -.05 (.06) | F(1, 279) = 2.88 | .09 | .01 |

Abbreviations: L, left; R, right. *Please see Figures 2C-E for pairwise comparisons. Grey shading indicates significant differences between trajectory means.

**Table S2**. Characterization of amygdala model-derived cortisol trajectories

| | Trajectory mean (SD) | | | | | | |
|---|---|---|---|---|---|---|---|
| | Hyper-responder | Elevated baseline | Non-responder | Responder | Statistic(df) | p | Effect size |
| Age | 23.56 (± 3.68) | 27.42 (± 6.76) | 25.31 (± 6.82) | 25.20 (± 8.25) | F(3, 277) = 0.61 | .60 | η² = .01 |
| Sex | | | | | χ²(3) = 33.31 | < .01 | V = .34 |
| M | 18 | 10 | 36 | 72 | | | |
| F | 0 | 2 | 29 | 114 | | | |
| BDI-II | 11.78 (±12.60) | 12.58 (±10.57) | 11.17 (±9.71) | 14.25 (±12.07) | F(3, 271) = 1.27 | .28 | η² = .01 |
| TICS | 15.39 (±9.62) | 15.92 (±10.12) | 16.37 (±8.85) | 18.53 (± 9.60) | F(3, 275) = 1.41 | .24 | η² = .02 |
| ASI (Total) | 14.59 (±6.73) | 17.00 (±9.30) | 18.94 (±11.21) | 20.95 (±10.98) | F(3, 272) = 2.41 | .06 | η² = .03 |
| CTQ (Total) | 31.06 (±7.49) | 35.33 (±10.66) | 31.48 (±5.04) | 32.41 (±7.95) | F(3, 273) = 1.09 | .35 | η² = .01 |
| LEC (Total) | 69.94 (±6.38) | 68.00 (±9.54) | 66.88 (±10.66) | 67.08 (±13.53) | F(3, 277) = 0.33 | .80 | η² = .00 |

Abbreviations: ASI = Anxiety Sensitivity Index, BDI-II = Beck Depression Inventory II, CTQ = Childhood Trauma Questionnaire, LEC = Life Events Checklist, TICS = Trier Inventory for the Assessment of Chronic Stress (Screening Scale). Grey shading indicates significant differences between trajectory means.

**Table S3.** Mean hippocampus predictor values across hippocampus-derived cortisol trajectories

| | Trajectory mean (SD) | | | | | |
|---|---|---|---|---|---|---|
| | Hyper-responder | Elevated baseline | Mild-responder | Statistic(df) | *p* | *η²* |
| Volume (mm³) | | | | | | |
| L | 4256.93 (350.33) | 4241.36 (252.12) | 3943.44 (366.57) | F(1, 279) = 13.58 | < .01* | .05 |
| R | 4262.74 (285.68) | 4313.63 (225.90) | 4028.26 (375.54) | F(1, 279) = 8.76 | < .01* | .03 |
| Activity (z) | | | | | | |
| L | -.03 (.03) | .02 (.02) | -.02 (.03) | F(1, 278) = 1.02 | .31 | .00 |
| R | -.02 (.02) | .03 (.02) | -.02 (.03) | F(1, 278) = 4.56 | .03* | .02 |
| Slope (z) | | | | | | |
| L | -.06 (.03) | .00 (.05) | -.04 (.05) | F(1, 278) = .06 | .80 | .00 |
| R | -.04 (.03) | -.00 (.04) | -.04 (.05) | F(1, 277) = .55 | .46 | .00 |

Abbreviations: L, left; R, right. *Please see Figures 3C-E for pairwise comparisons. Grey shading indicates significant differences between trajectory means.

**Exploratory affect analyses**

First, we explored whether cortisol trajectories from amygdala and hippocampus models differed in negative affect. A mixed-effects ANOVA on PANAS negative affect using cortisol groupings from the four-trajectory amygdala model revealed no significant differences between trajectories (main effect of time: $F(3.79, 963.27) = 45.44$, $p < .00$, $\eta_p^2 = .15$, main effect trajectory: $F(3.00, 254.00) = 1.31$,

$p = .27$, $\eta_p^2 = .02$, time*trajectory interaction: $F(11.38, 963.27) = .82,\ p = .62,\ \eta_p^2 = .08)$. See Figure S2A for a visualization. A mixed-effects ANOVA on negative affect with cortisol-derived trajectories from the hippocampus model also gave no indication for differential affect across cortisol trajectories (main effect time: $F(3.77, 961.95) = 18.66, p < .00, \eta_p^2 = .07$, main effect trajectory: $F(2.00, 255.00) = .35,\ p = .71,\ \eta_p^2 = .00$, time*trajectory interaction: $F(7.55, 961.95) = 1.12,\ p = .35,\ \eta_p^2 = .01$, Figure S2B).

When cortisol trajectories did not show utility in explaining variance in negative affect, we further explored whether re-running baseline and brain models with negative affect as the outcome may facilitate the identification of distinguishable affect trajectories (Table S1). Among baseline models, all fit indices unanimously (BIC, SABIC, AIC) selected the three-trajectory model with low entropy $(.48)$. The first trajectory exhibited 'strong-lasting' negative affect in response to stress $(N = 78)$, the second trajectory had a moderate-brief negative affective response $(N = 100)$, and the final trajectory had a mild-brief negative response $(N = 101)$. The baseline affect model is displayed in Figure S4A.

In the following, we compared this winning baseline model to brain models. For amygdala, hippocampus, and rostral anterior cingulate cortex (rACC) models, BIC, SABIC, and entropy indices (all .81) favored two-trajectory models, while AIC favored the three-trajectory models. Hence, we explored both. The two-trajectory models of all three ROIs converged on similar points (Table S1), leading to identical trajectory assignment across models $(Fleiss'\ kappa = 1,\ z = 28.9,\ p < .01)$. Descriptively, the first trajectory showed a 'strong negative affect' trajectory $(N = 71)$ and the second a 'moderate negative affect' trajectory $(N = 208)$. Posterior polynomial logistic regression indicated that brain parameters of the three ROIs showed no utility in predicting membership of two-trajectory affect models $(all\ p's > .05)$. The same was true for four-trajectory models. Convergence points of the three ROIs were similar (Table S1) and so were trajectory assignments $(Fleiss'\ kappa = .89,\ z = 42.2,\ p < .01)$. Descriptively, the four trajectories followed similar patterns, but varied in intensity, from 'strong-lasting' negative affect in the first trajectory towards 'mild-brief' negative affect in the fourth trajectory. The first trajectory comprised the smallest sample portion $(N = 28)$, while trajectories two to four had approximately equal proportions, with $N = 80, 83, 88$, respectively. Additionally, none of the ROIs predicted the four negative affect trajectories in posterior analyses $(all\ p's > .05)$. We therefore only display amygdala two- and four-trajectory models (Figure S3B&C). However, due to the similarity between models, amygdala affect models may also be considered as stand-ins for hippocampus and rACC models.

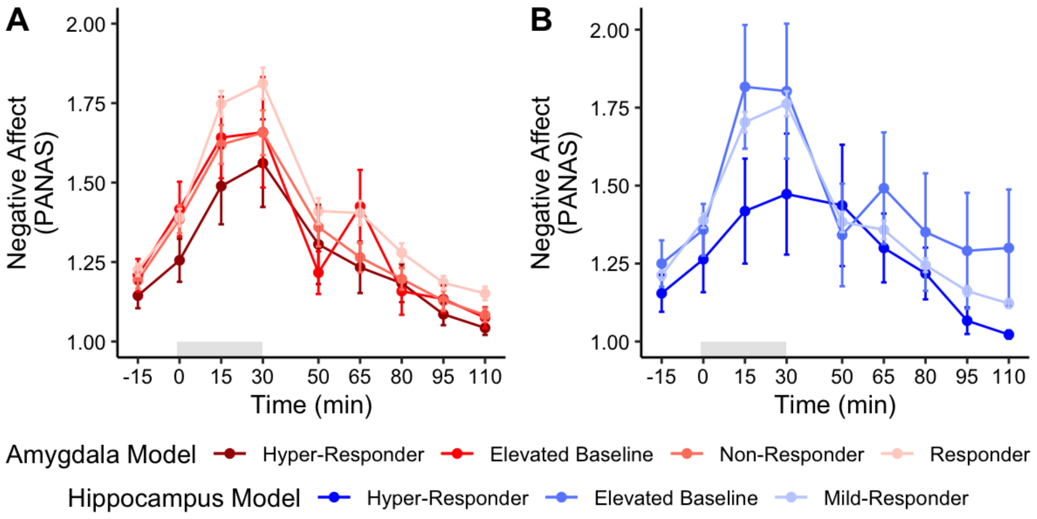

**Figure S2.** Negative affect across cortisol trajectories (A) from the 4-trajectory amygdala model and (B) the 3-trajectory hippocampus model. PANAS, Positive and Negative Affect Schedule. Grey shading indicates the timing of Scan*STRESS*.

**Table S4**. Between model comparison of negative affect models

| Model | Loglik[1] | Npm[2] | AIC | BIC | SABIC | Entropy | Trajectory size (%) | | | |
|---|---|---|---|---|---|---|---|---|---|---|
| | | | | | | | 1 | 2 | 3 | 4 |
| **Baseline models** | | | | | | | | | | |
| 1-Trajectory | -86.76 | 13 | 199.53 | 246.73 | 205.51 | **1.00** | 100 | NA | NA | NA |
| 2-Trajectory | -83.92 | 16 | 199.85 | 199.85 | 257.95 | .44 | 63.08 | 36.91 | NA | NA |
| 3-Trajectory | -54.02 | 19 | **146.05** | **215.04** | **215.04** | .48 | 27.95 | 35.84 | 36.20 | NA |
| 4-Trajectory | -54.02 | 22 | 152.04 | 231.93 | 231.93 | .41 | 28.31 | 48.74 | .00 | 22.93 |
| **Amygdala models** | | | | | | | | | | |
| 2-Trajectory | -53.37 | 19 | 144.75 | **213.74** | **153.49** | **.81** | 25.44 | 74.55 | NA | NA |
| 3-Trajectory | -52.06 | 25 | 154.13 | 244.91 | 165.63 | .51 | 45.16 | 27.24 | 27.59 | NA |
| 4-Trajectory | -39.39 | 31 | **140.79** | 253.35 | 155.06 | .67 | 10.03 | 28.67 | 29.74 | 31.54 |
| **Hippocampus models** | | | | | | | | | | |
| 2-Trajectory | -53.29 | 19 | 144.58 | **213.57** | **153.33** | **.81** | 25.44 | 74.55 | NA | NA |
| 3-Trajectory | -51.71 | 25 | 153.42 | 244.20 | 164.93 | .51 | 27.24 | 21.50 | 51.25 | NA |
| 4-Trajectory | -38.87 | 31 | 139.75 | 252.32 | 154.02 | .64 | 8.60 | 29.39 | 29.03 | 32.97 |
| **mOFC models** | | | | | | | | | | |
| 2-Trajectory | -54.07 | 19 | 146.14 | 215.14 | 154.89 | **.81** | 25.44 | 74.55 | NA | NA |
| 3-Trajectory | -51.75 | 25 | 153.51 | 244.29 | 165.02 | .54 | 27.24 | 45.51 | 27.24 | NA |
| 4-Trajectory | -34.40 | 31 | **130.81** | 243.38 | **145.08** | .67 | 11.11 | 25.08 | 24.73 | 39.06 |
| **rACC models** | | | | | | | | | | |
| 2-Trajectory | -53.12 | 19 | 144.25 | **213.25** | **153.00** | **.81** | 25.44 | 74.55 | NA | NA |
| 3-Trajectory | -48.96 | 25 | 147.92 | 238.70 | 159.43 | .62 | 31.89 | 41.21 | 26.88 | NA |
| 4-Trajectory | -38.41 | 31 | **138.83** | 251.40 | 153.10 | .66 | 11.46 | 25.80 | 31.89 | 30.82 |

[1]Log likelihood, [2]Number of parameters, grey shading indicates selected models.

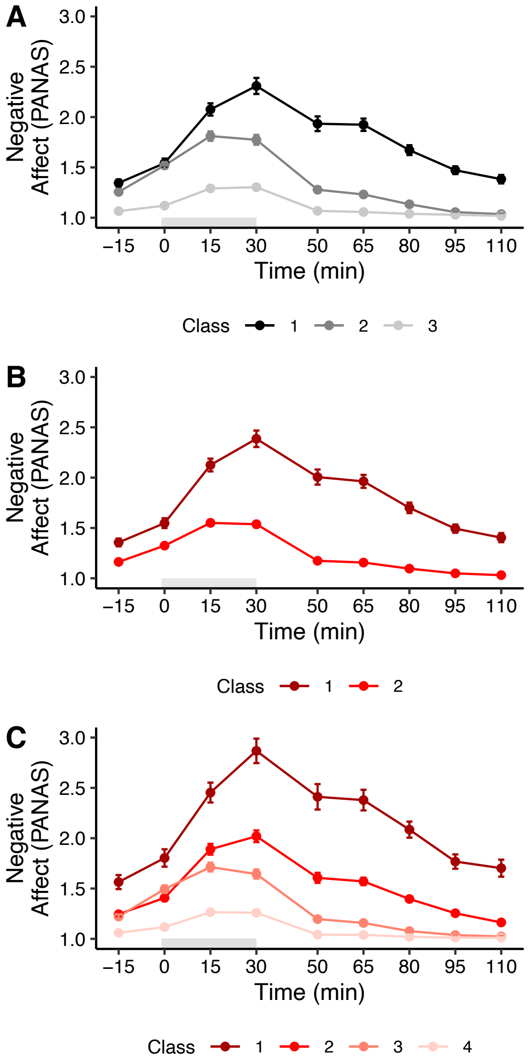

**Figure S3.** Baseline and amygdala negative affect models. A) Best fitting baseline model without additional brain predictors. B + C) Amygdala-derived two- and four-trajectory negative affect models. Note that amygdala plots may be read as surrogates for rACC and hippocampus affect models, which had nearly identical trajectories and assignments. Grey shading indicates Scan*STRESS* timing.

Furthermore, we explored whether cortisol trajectories from the selected amygdala and hippocampus models corresponded with affect models with the same trajectory numbers, as well as the winning two-trajectory affect model. There was no meaningful correspondence between cortisol and affect trajectory assignment for neither amygdala nor hippocampus models. See Figure S4 for an example of such non-correspondence in amygdala models.

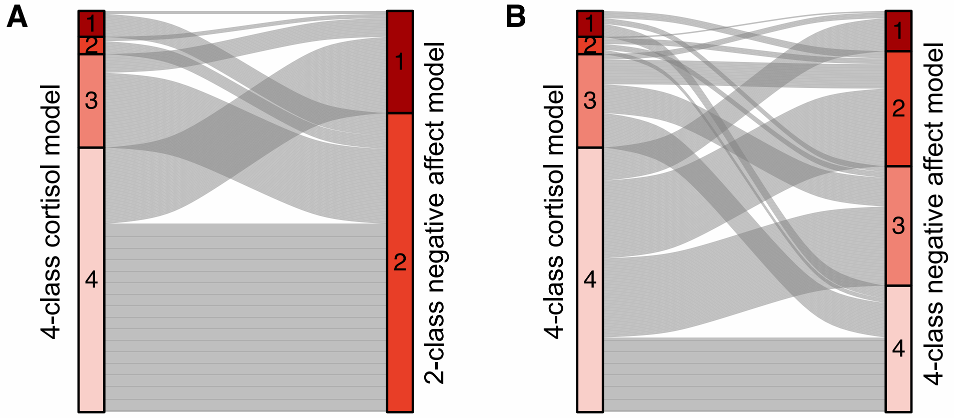

**Figure S4.** Alluvial plots depicting participant flow between amygdala cortisol and negative affect models. A) Best-fitting four-trajectory cortisol model and two-trajectory negative affect model. B) Best-fitting four-trajectory cortisol model and four-trajectory negative affect model.
